# EMC regulates cell membrane fluidity to facilitate biogenesis of membrane proteins

**DOI:** 10.1101/2024.10.15.618411

**Authors:** Modesto Berraquero, Víctor A. Tallada, Juan Jimenez

## Abstract

The EMC complex is a highly conserved transmembrane chaperone located in the endoplasmic reticulum (ER). This complex has been associated in humans with sterol homeostasis and a myriad of different cellular activities, making the mechanism of EMC functionality enigmatic. Using fission yeast, we demonstrate that EMC assists biogenesis of the sterol transfer protein Lam6/Ltc1 at the ER-Plasma membrane and ER-Mitochondria contact sites. Cells losing EMC function sequester unfolded Lam6/Ltc1 and other proteins at the mitochondrial matrix, driving to surplus ergosterol, cold-sensitive growth and mitochondrial disfunctions. Remarkably, inhibition of the ergosterol biosynthesis, but also fluidization of cell membranes to counteract its rigidizing effects, reduce the ER-unfolded protein response and rescue growth and mitochondrial defects in EMC-deficient cells. These results indicate that EMC-assisted biogenesis of Lam6/Ltc1 provides, through ergosterol homeostasis, optimal membrane fluidity to facilitate biogenesis of other ER-membrane proteins.

**HIGHLIGHTS:**

- Lam6/Ltc1 requires EMC for localization at ER-Plasma membrane and ER-Mitochondria contact sites.
- EMC-driven biogenesis of Lam6/Ltc1 is involved in membrane fluidity homeostasis.
- EMC-deficient cells accumulate misfolded proteins in the mitochondrial matrix.
- Cell membrane fluidization alleviates the ER-misfolded proteins response and rescues growth and mitochondrial defects of EMC-deficient cells.

**Graphical Abstract:** 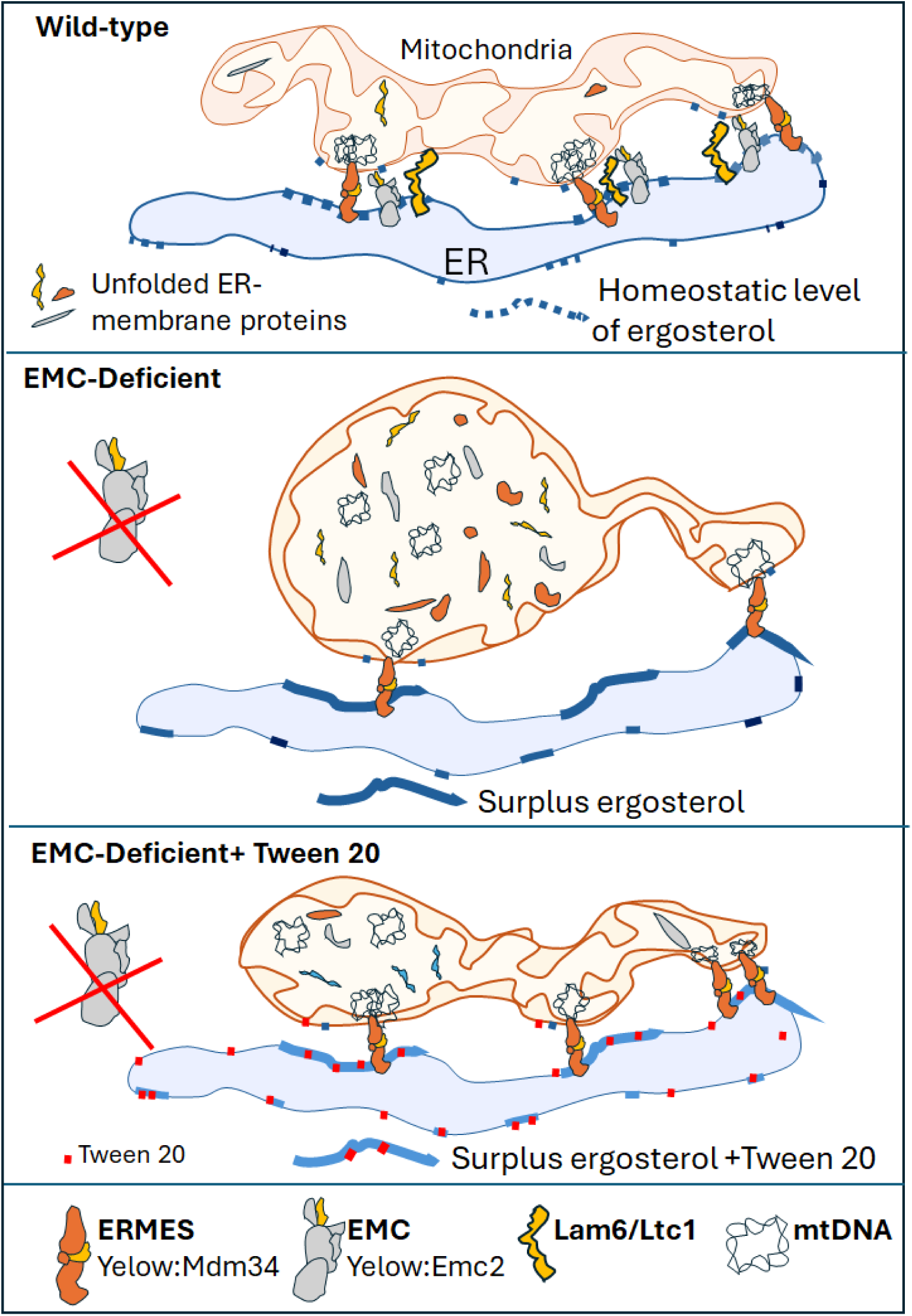

Schematic representation of EMC-membrane fluidity homeostasis connecting protein folding and mitochondrial activity.

## INTRODUCTION

The endoplasmic reticulum (ER) is a vast multifarious membrane system in eukaryotic cells, the primary site for lipids and carbohydrates biosynthesis, as well as the folding, assembly, modification, and transport of secreted and integrated membrane proteins.^1,2^

It is known to house many protein complexes that aid its function, including the conserved ER membrane protein complex (EMC).^3^

The EMC was first described in the budding yeast *Saccharomyces cerevisiae* as a complex composed of 6 proteins with similar genetic interaction patterns^1^. The structural integrity of the EMC depends on ‘core subunits’ EMC1, EMC2, EMC3, EMC5 and EMC6, which are essential for both assembly and function of the complex.^4–6^ Two other membrane proteins, EMC7 and EMC10, also copurify with this complex to yield a mature eight-subunits complex.^7^ An MS-based mapping of the ER-associated degradation (ERAD) interaction network not only confirmed the EMC orthologous in mammals but also identified EMC8 and EMC9, two metazoan-specific subunits.^8^

Biogenesis of membrane proteins et the ER requires insertion into lipid bilayers with specific topologies to carry out their functions.^1,4^ The EMC assists many of them by promoting folding and insertion of their atypical and sub-optimal transmembrane domains (TMDs). Therefore, the EMC can function at the ER both co-translationally as a chaperone for folding TMD-containing proteins and post-translationally as an insertase for tail-anchored proteins.^9^

Notably, dysfunction of the EMC complex has wide-ranging consequences such as ER stress, lipids transport and homeostasis, and organelle tethering, among others.^3^ Unravelling the primary relationship between EMC and the multifaceted cellular consequences of altering its client proteins remains a formidable challenge.

The fission yeast *Schizosaccharomyces pombe* is an important single-cell model organism widely used to study various aspects of eukaryotic cellular and molecular biology. In a genome-wide search for genes causing overexpression cell cycle arrest in *S. pombe* cells (“*oca*” genes), we previously identified the *oca3* gene^10^, which was found to encode the putative EMC2 orthologous in vertebrates (Pombase ID SPBC15C4.01c)^11^. By analyzing the consequences of Oca3/Emc2 depletion, here we asked for the cellular and molecular mechanisms connecting EMC and the pleiotropic phenotypes observed in EMC dysfunctional *S. pombe* cells. Our results uncover that membrane fluidity homeostasis regulated by EMC is on the root of these cell defects.

## RESULTS

### Oca3 encodes the fission yeast ortholog to the metazoan EMC2

Sequence similarity predicts that the *oca3* gene encodes the putative fission yeast ortholog for the EMC2 subunit of the eukaryotic EMC complex.^12^ To experimentally support this prediction, the encoded Oca3 protein was tagged with mCherry and expressed from its native locus in *S. pombe* cells co-expressing the ADEL-GFP fusion, a bona-fide ER marker.^13^ As expected for an EMC component, *in vivo* fluorescence microscopy shows that Oca3-mCherry localizes to the ER (Fig. 1A), co-localizing with the ER-reporter ADEL-GFP. Localization of the Tts1-mCherry construct, enriched in the tubular structure of the ER^14^ indicates that the ER, ER-associated plasma membrane (PM) and nuclear membrane remain unaltered in *oca3*-null cells (the *oca3Δ* deletion strain) (Fig. 1B).

**Fig. 1.**
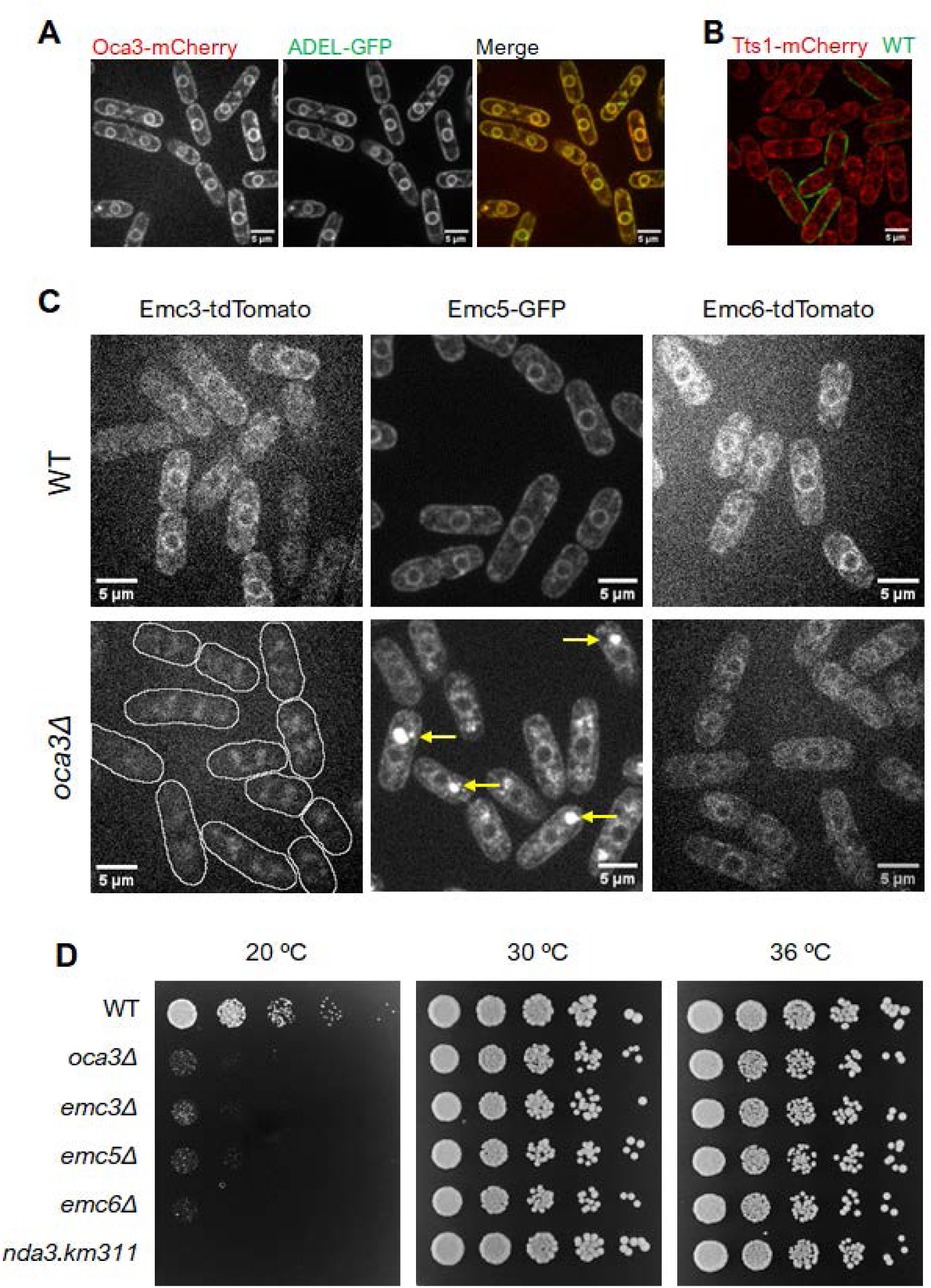
Genetics analysis of EMC complex proteins. **A**. Subcellular localization Oca3-mCherry (left panel) and the ER reporter ADEL-GFP (central panel). Merge image is shown (right panel). **B**. Membrane organization of *oca3Δ* mutant vs *wild-type* cells. Fluorescent images of Tts1-mCherry (protein located in most subcellular membranes; control *wild-type* cells are stained with green lectin) **C**. Subcellular localization of Emc3-tdTmato, Emc5-GFP and Emc6-mCherry subunits of the EMC complex in *wild-type* and *oca3Δ* genetics background. In *oca3Δ* cells, Emc3 is missing from the ER (lines are used to shape cells), Emc5 forms aggregates (arrows) and Emc6 decreases fluorescence intensity. **D**. Spot growth test (sequential five-fold dilutions) of *oca3Δ*, *emc3, emc5Δ,* and *emc6Δ* strains, along with *wild-type* (wt) and *nda3.km311* as a cold-sensitive control at 20 °C, 30 °C and 36°. Scale bar 5µm.

In budding yeasts, the eight subunits of the EMC heterooligomer complex show similar genetic interaction patterns.^1^ The cytosolic region of EMC is formed by Emc2 and the cytosolic domains of Emc3, Emc4 and Emc5, whereas Emc1 and Emc3–Emc6 are transmembrane proteins.^15^ Like Oca3, fluorescently tagged constructs of the putative orthologous *S. pombe* subunits Emc3 (Pombase ID SPBC1711.03), Emc5 (Pombase ID SPAP4C9.02) and Emc6 (Pombase ID SPCC1020.11c) localize to the ER (Fig. 1C). As shown in Fig. 1C, depletion of Oca3 leads to loss of Emc3, aggregation of Emc5, and normal ER localization, but reduced fluorescence levels, of EMC6, revealing the requirement of Oca3 for the assembly of a functional EMC complex.

The *oca3Δ* strain yields sick cells that can grow at 30°C but are unable to grow at lower temperatures (routinely assessed at 20°C) (Fig. 1D), resulting in a cold-sensitive growth phenotype. Likewise, the deletion of genes encoding the above predicted components of the *S. pombe* EMC (*emc3Δ*, *emc5Δ* and *emc6Δ*) renders a cold-sensitive phenotype too (Fig. 1D). Taken together, we conclude that the *oca3* gene encodes the eukaryotic EMC2 ortholog found in *S. pombe* (Oca3/Emc2), the deletion of this gene driving to EMC-deficient cells.

### EMC deficiency leads to mitochondrial dysfunction

Depletion of Oca3/Emc2 leads to severe growth defects at low temperature. To determine possible roles of EMC in *S. pombe* cell growth, we first observed nuclear and mitochondrial DNA in DAPI-stained cells under permissive (30°C) and restrictive (20°C) growth conditions. Compared to the *wild-type* control, no obvious alterations in nuclear staining can be observed in *oca3Δ* cells, but clumps of mitochondrial DNA (mtDNA) are observed in the cytoplasm of these mutant cells (Fig. 2A). Quantitative PCR analysis of target mtDNA sequences determined a 28% reduction in mtDNA molecules per cell in the mutant strain at 30°C, exacerbated at low temperature (37% reduction at 20°C) (Fig. 2B). This result suggests that EMC may be required for the proper organization and/or stability of mtDNA during its replication/segregation cycle.

**Fig. 2.**
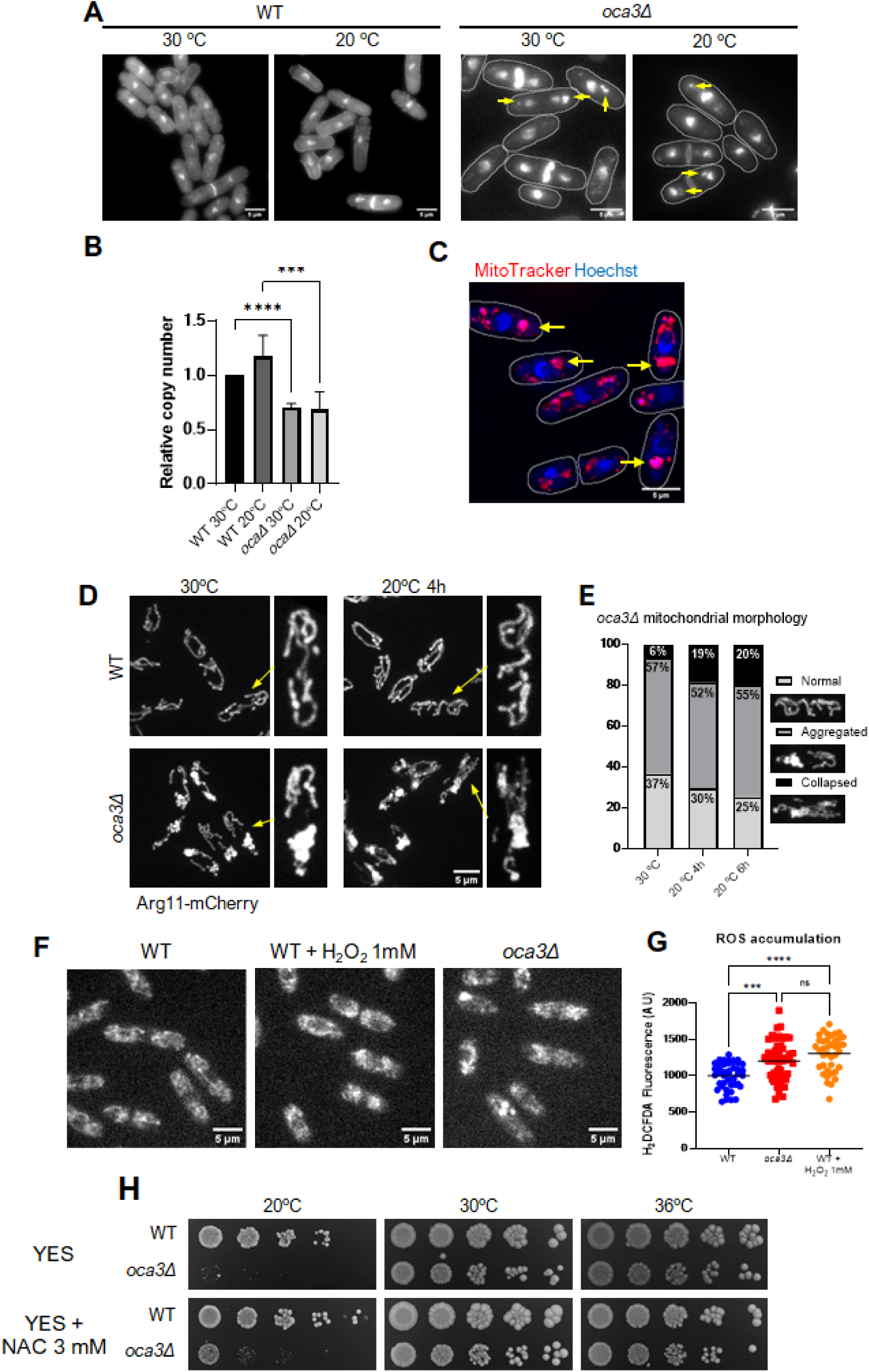
EMC dysfunction alters mitochondrial structure and function. **A.** DNA visualisation via DAPI staining both at permissive (30°C) and restrictive (20°C after 6h incubation) temperatures. Arrows indicate DNA aggregates outside the nucleus. Images are max projections of stacks of 0.5 µm layers. **B.** mtDNA copy number/cell quantified by qPCR in *oca3Δ* and *wild-type* (wt) cells at 30°C and 20°C. Statistical significance is indicated (number of *). **C.** Fluorescence images obtained from DNA (Hoechst) and mitochondria (MitoTracker) staining of Oca3-depleted cells at 30°C. Mitochondrial membranes and mtDNA aggregations co-localize (arrows; lines indicating cell shape). **D.** *in vivo* Arg11-mCherry localization (mitochondrial lumen marker) in *oca3Δ* (lower panels) and the *wild-type* control (upper panels) cells at 20°C (right panels) and 30°C (left panels). Deletion of *oca3* condensates the mitochondrial structure resulting in viable mitochondrial aggregation at 30 °C, exacerbated after 4h at 20°C where the mitochondrial network collapses (arrows and enlargements). **E.** Quantification of mitochondrial Arg11-mCherry fluorescence (see methods). Representative cells with either normal, aggregated and or collapsed mitochondria structures are shown. High proportion of collapsed mitochondrial membranes after 4h (19%, n=176) and 6h (20%, n=95) incubation at 20°C (restrictive growth temperature), as compared to cells incubated at 30°C (permissive growth temperature) (6%, n=93), is observed. Scale bars 5µm. **F.** H_2_DCFDA-stained (ROS accumulation indicator) *wild-type* (wt, n=44) and *oca3Δ* (n=53) cells at 30°C (microphotographs) to measure the accumulation of reactive oxygen species (ROS). Incubation of *wild-type* cells with 1mM de H_2_O_2_ (n=38) was used as a positive control Scale bars 5µm. **G.** Fluorescence quantification of H2DCFDA was used to estimate ROS levels. **H.** Spot growth assay of *oca3Δ* and *wild-type* (wt) cells in rich media (YES) with and without 3mM N-Acetylcysteine (NAC) at 20°C, 30°C and 36°C (\*\*\**p* value <0.005, \*\*\*\**p* value <0.001; ns: not significant)

To determine whether mtDNA aggregations generated in *oca3Δ* cells are accompanied by alterations in the mitochondrial structure, mtDNA and mitochondrial membranes were stained using Hoechst and MitoTracker respectively and cells were analysed under the microscope. As shown in Fig. 2C, Oca3/Emc2 depletion results in a condensed mitochondrial structure that co-localize with abnormal mtDNA aggregations, revealing severe mitochondrial dysfunctions (Fig. 2D-E) in EMC-deficient cells.

The accumulation of reactive oxygen species (ROS) is associated to mitochondrial dysfunctions.^16^ As shown in Fig. 2F-G, proliferating Oca3-depleted cells (at 30°C) reach ROS levels similar to those found in *wild-type* cells subjected to oxidative stress (H_2_O_2_ treatment), indicating that EMC-deficient cells suffer mitochondrial stress. The addition of N-Acetylcysteine (NAC), a compound known to act as a reducing agent and protect cells from excess ROS^17^, slightly restores growth of *oca3Δ* cells at 20°C (Fig. 2H), suggesting that mitochondrial dysfunction, among other reasons, is likely compromising the survival of these EMC-deficient cells. Overall, we conclude that EMC is required for mtDNA stability and normal mitochondria structure and function.

### Oca3/Emc2 depletion impairs ER-Mitochondrial tethering

ER membranes show normal structure in *oca3Δ* cells (see in Fig 1). Therefore, mitochondrial aggregates observed in these mutant cells suggest that mitochondrial membranes are largely detached from ER membranes. Interestingly, the defective mitochondrial structure observed in *oca3*Δ mutant cells (Fig. 2D-E) resembles that reported in mutants with reduced ER-Mitochondrial tethering^18^, suggesting also a role for EMC in ER-Mitochondria tethering in *S. pombe*, as reported in mammals and budding yeasts.^19,20^ However, EMC anchoring functions could be exerted indirectly, by assisting the biogenesis of ER-Mitochondria docking factors such as ERMES (ER–Mitochondria Encounter Structure)^21^ or Lam6/Ltc1, a TMD-protein that enables sterols trafficking at ER-PM and ER-vesicles contacts in *S. pombe* cells^22^ and the ER-Mitochondria and ER– vacuole contact sites in *S. cerevisiae.*^23^

To further study the possibility that tethering complexes could be included in the wide range of putative EMC clients,^24^ the ERMES complex subunit Mdm34^25^ and Ltc1^22^ were tagged with GFP, and their subcellular localization was analysed by time-lapse microscopy in *wild-type* and *oca3Δ* cells at 30°C and 20°C.

As shown in Fig. 3, Mdm34-GFP co-localizes with mitochondrial membrane markers in *wild-type* cells, enriched at the ERMES ER-Mitochondrial contact sites.^26^ In *oca3Δ* cells, most of the Mdm34-GFP is delocalised into the abnormal mitochondria, indicating that EMC may facilitates ERMES biogenesis. But eventually, patches of Mdm34-GFP are still observed in the outer membrane in the remaining mitochondrial tubular structures of this mutant (see in Fig. 3), suggesting that EMC is not essential for ERMES assembly. Loss of ERMES function is lethal in *S. pombe*^18^, suggesting that the fraction of this complex found at ER-Mitochondria contacts is still functional (see Fig. 3).

**Fig. 3.**
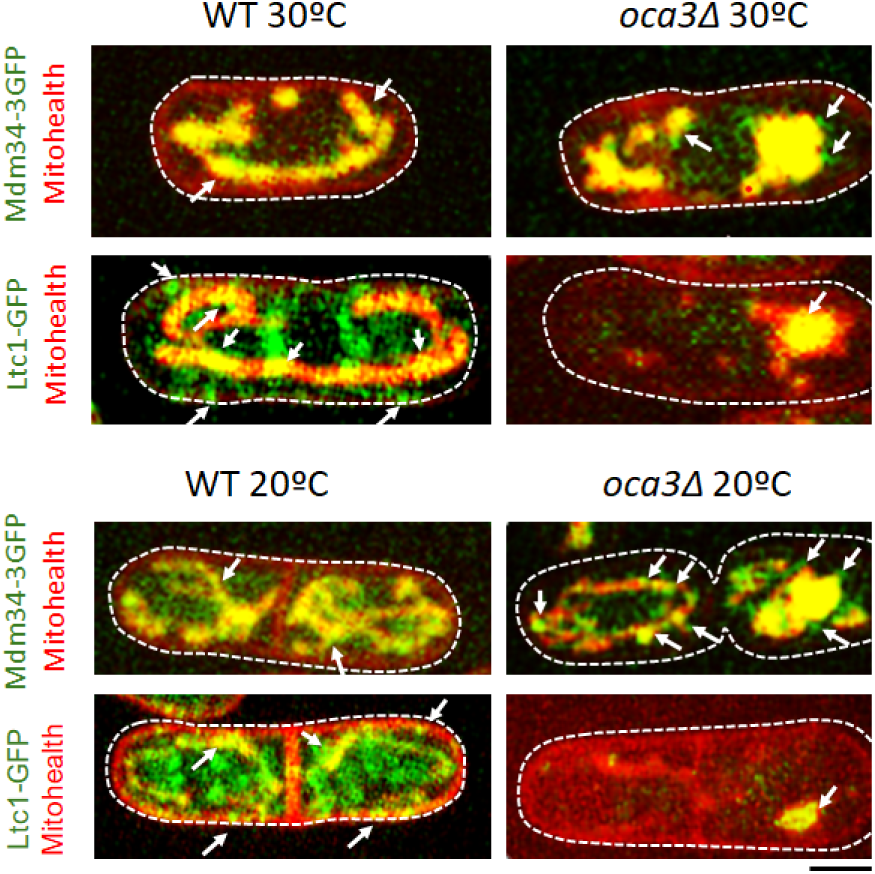
ERMES and Ltc1 subcellular localization. Fluorescence images of the ERMES subunit Mdm34-3GFP and Ltc1-GFP in *wild-type* (wt) and *oca3Δ* representative cells at 30°C (top panels) and 20ªC (lower panels). Mitohealth (red) and GFP (green) fluorescence are used in the same cells (merge images) to highlights mitochondrial and PM membrane localization and co-localization (yellow). Scale bar 5 µm.

In agreement with its localization in ER-PM and ER-vesicles previously described in fission yeast^22^, Ltc1-GFP localizes to cortical and internal dots in the ER in *wild-type* cells (arrows in Fig. 3), but Mitohealth co-localization indicates that this protein is also associated to ER-Mitochondrial contact sites in *S. pombe* cells (Fig. 3 and Suppl. Fig. S1), as reported in *S. cerevisiae* cells.^23^ In contrast to Mdm34-GFP, Ltc1-GFP lacks its normal ER-PM and ER-Mitochondria localization (see Fig. 3). This result indicates that Ltc1 biogenesis is fully dependent on EMC activity.

Interestingly, Ltc1-GFP delocalises from the ER-membrane contact sites to decorate the matrix of the mitochondrial aggregates in *oca3*Δ cells (see in 3). The sequestration of Ltc1-GFP into the mitochondria in these cells can be attributed to the mitochondria-associated proteostatic mechanism for unfolded proteins in the ER (ER-associated mitochondrial sequestration, ERAMS).^27^ We therefore conclude that Ltc1 is a client of EMC, whose proper folding and biogenesis in the ER is driven by the EMC. Overall, our results suggest that EMC may be involved in ER-Mitochondrial tethering by favouring folding and insertion of ERMES and by assisting Ltc1 biogenesis.

### EMC regulates ergosterol homeostasis by assisting Ltc1 biogenesis

The EMC complex has emerged as an important eukaryotic complex for homeostasis of sterols ^6,9^. Since *S. pombe ltc1Δ* mutant cells lead to the overaccumulation of ergosterol in yeast cell membranes ^22^, we wondered whether Ltc1 delocalization (Fig. 3) also resulted in overaccumulation of ergosterol in the *oca3Δ* strain.

To determine whether EMC is involved in ergosterol homeostasis in *S. pombe* cells, sterols were extracted from cells incubated at 20°C and 30°C and the amount of ergosterol was quantified by TLC silica gel. As shown in Fig. 4A, loss of EMC drives to a marked increase in the ergosterol content in the *oca3Δ* mutant strain at both temperatures when compared to levels in *wild-type* cells. Increased levels are also observed by fluorescent quantification of sterol droplets using the D4H probe^22^ (Fig. 4B). Physiological assays were also used to confirm these analytical results. Nystatin specifically binds ergosterol in the yeast PM creating pore-forming complexes that result in cell death.^28^ Consequently, mutants that reduce ergosterol content are particularly resistant to the effects of nystatin^29^, whereas high ergosterol levels render cells more sensitive to the effects of the drug.^30^ On the contrary, cells that accumulate excess ergosterol are highly resistance to ketoconazole, a drug that reduces ergosterol content by inhibiting its biosynthesis.^31^ As expected for the observed ergosterol overaccumulation (Fig. 4A-B), the growth of Oca3/Emc2-depleted cells is sensitive to nystatin and resistant to ketoconazole (Fig. 4C). Since Ltc1 requires EMC for its localization and function at ER-Plasma membrane and ER-Mitochondria contact sites (Fig. 3), we conclude that EMC regulates ergosterol homeostasis through the ER-assisted biogenesis of the sterol transfer protein Ltc1.

**Fig. 4.**
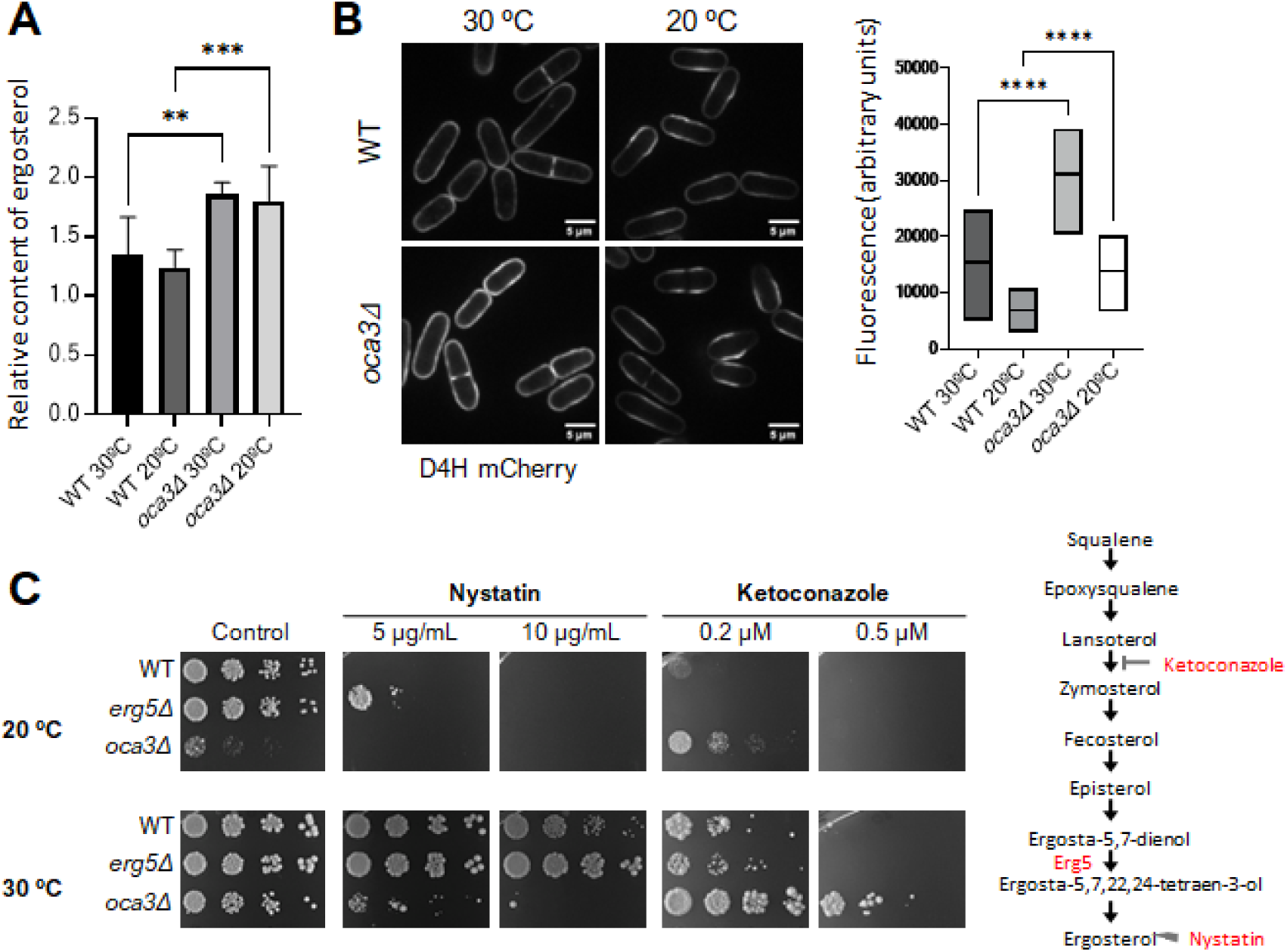
Overaccumulation of ergosterol in the *oca3Δ* mutant strain. **A** Quantification of ergosterol content by TLC analysis in *oca3Δ* and *wild-type* (wt) control cells at 20°C and 30°C (statistical significance is indicated (\*\**p* value <0.01, \*\*\**p* value <0.005). **B.** Fluorescence images of D4H probe in *wild-type* (wt) and *oca3Δ* cells at 20°C and 30°C with its quantification (\*\*\*\**p* value <0.001). N>40, scale bar 5 µm. **C.** Spot growth assay (sequential five-fold dilutions) of *wild-type* (control) and *oca3Δ* strains at 20ªC and 30°C on control plates and plates containing nystatin and ketoconazole. The *erg5Δ* strain, which disrupts the ergosterol synthesis^29^ is used as control. Steps of the ergosterol biosynthesis affected by *erg5Δ,* ketoconazole and nystatin are highlighted.

Strikingly, ketoconazole rescues the cold-sensitive growth phenotype of Oca3/EMC2-depleted cells (see in Fig. 4C), indicating that ergosterol overaccumulation is indeed a major cause of the detrimental growth of these *oca3Δ* cells. Equivalent results were reported in mammals, where the excess cholesterol (the equivalent to ergosterol in mammals) in EMC-depleted mutants leads to cell death.^6^

### Membrane fluidization rescues cell functions in EMC-deficient cells

The cold-sensitive growth rescued by ketoconazole indicates that the surplus ergosterol largely accounts for growth defects of Oca2/Emc2-depleted cells at low temperature (20°C) (see Fig. 4C). Ergosterol (cholesterol in higher eukaryotes) is a cell membrane-rigidizing molecule^32,33^, its homeostatic regulation being necessary to maintain optimal lipids bilayer fluidity under environmental changes.^34–37^ Therefore, we hypothesised that upon ergosterol overaccumulation, membrane fluidity in *oca3Δ* cells is below the survival threshold at 20°C. The increase in fluidity with temperature may counteract this rigidifying effect thereby providing a permissive growth condition for this mutant strain at high temperature (assessed at 30°C and 36°C) (Fig. 1D, Fig. 2H and Fig. 4C). To further investigate this idea, we analyzed the effects on growth of different membrane-fluidizing agents (detergents and alcohols)^38–40^ in *oca3Δ* cells. As shown in Fig. 5A, ethanol, a membrane fluidizer at intermediate concentrations^40,41^ partially suppresses sensitive growth at 20°C in this strain. Sorbitol shows similar results, but the non-ionic detergent tween 20 is surprisingly efficient in rescuing cold-sensitive growth of *oca3Δ* cells (Fig. 5B). At this temperature (20°), ergosterol content is low in *wild-type* cells, but increase in media with tween 20 (Fig. 5C), as expected for the homeostatic response to counteract the fluidizing effects of this solvent. EMC-deficient cell lacks ergosterol homeostasis, and ergosterol levels remain high regardless of tween 20 addition and growth temperature (20°C and 30°C) (Fig. 5C). Therefore, as temperature, tween 20 likely rescues cold-sensitive growth of EMC-deficient cells through its fluidizing properties, by counteracting membrane rigidizing of surplus ergosterol in these mutant cells, rather than by reducing its membrane content (as ketoconazole).

**Fig. 5.**
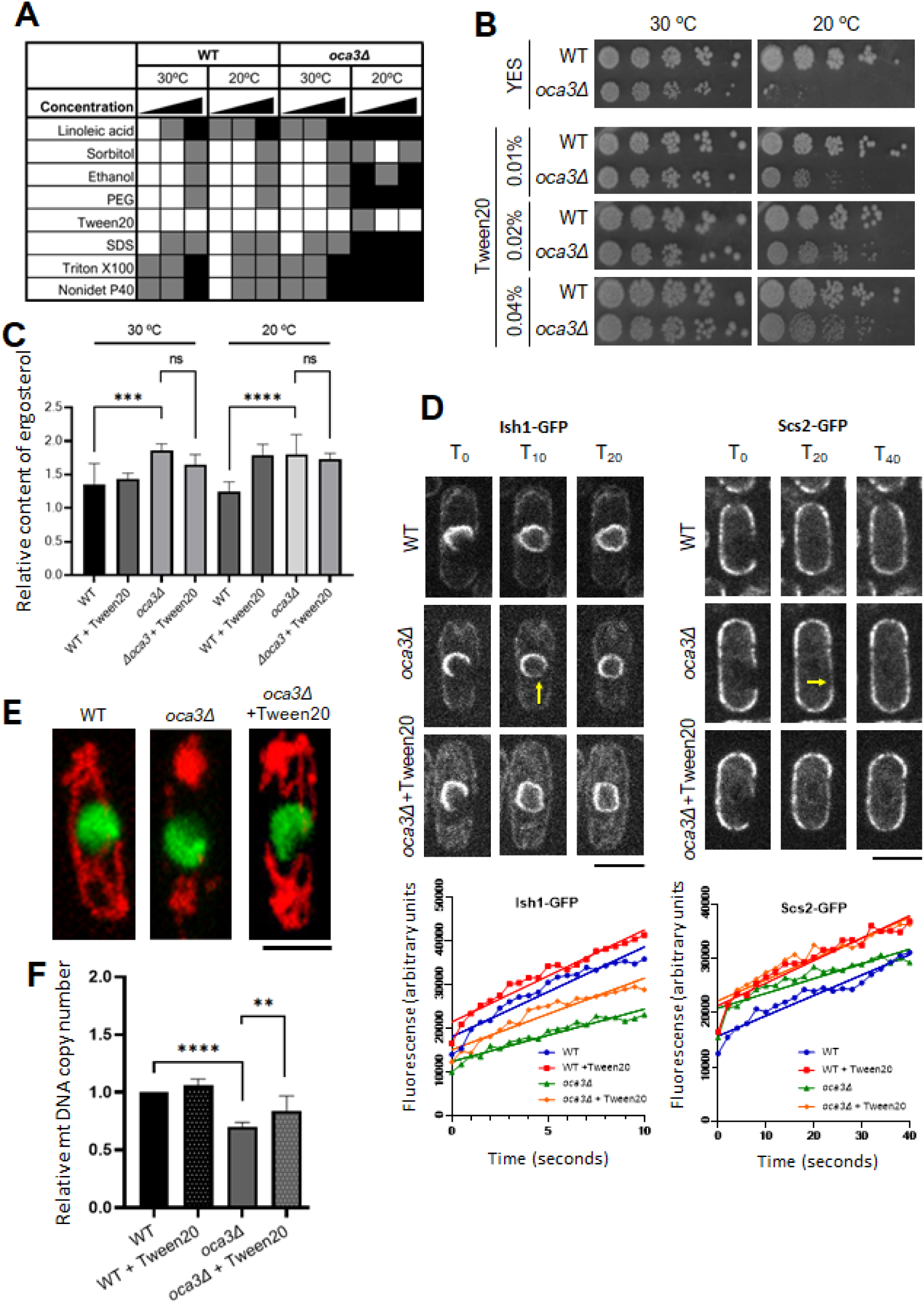
Chemical suppression of *oca3Δ* defects by membrane fluidifying compounds. **A.** Effects on growth of *oca3Δ* cells by the indicated chemicals at increasing concentrations. White indicate viability while black represents lethality. **B.** Spot growth assay of *wild-type* (wt) and *oca3Δ* cells at the indicated tween 20 concentration (%v/v) and temperature (20°C and 30°C) (control media YES is used). Tween 20 (0,02-0.04 % v/v) efficiently restores viability of *oca3Δ* cells at 20°C. **C.** Quantification of ergosterol content by TLC analysis in *wild-type* and *oca3Δ* cells in the presence of tween 20 (0.04%) at 20°C and 30°C (squared graph). Data from control experiments without this solvent are included (previously shown in Fig. 4A) (\*\*\**p* value <0.005, \*\*\*\**p* value <0.001, ns: not significant). **D.** Fluorescence recovery by lateral diffusion of the nuclear envelop inserted Ish-GFP after Photobleaching (left) and the plasma membrane/ER inserted Scs2-GFP protein (right) in *wild-type* and *oca3Δ* cells with and without tween 20 as indicated. Scale bars 5µm. Their fluorescent recovery dynamics (graphs) are shown. Slopes are statistically significative between *wild-type* and *oca3Δ* cells both in *ish1-*GFP (2048 vs 1196 p-value <0.0001) and *scs2-*GFP (374 vs 271 p-value=0.01) and not significative between *wild-type* and *oca3Δ* with tween20. **E.** Representative image of nuclear DNA (green) and the membrane mitochondrial network (red) of *wild-type*, *oca3Δ* cells and *oca3Δ* cells with tween 20 (0.04% v/v) at 20°C and 30°C. Reduced aggregations and the recovery of tubular membranes in the mitochondrial network of *oca3Δ* cells with tween 20 indicates a partial recovery of *wild-type* structures by this non-ionic detergent (arrows). **F.** mtDNA copy number/cell quantified by qPCR in cells of *the indicated strains* at 30°C in the presence of tween 20 (0.04%) or without this detergent. Statistical significance is indicated (\*\**p* value <0.01, \*\*\*\**p* value <0.001).

To estimate *in vivo* fluidizing effects of tween 20 on ergosterol enriched membranes of Oca3/Emc2-depleted cells, Fluorescent Recovery After Photobleaching (FRAP) assays were carried out to study lateral diffusion of Scs2-GFP (ER/PM protein)^42^, and Ish1-GFP (nuclear envelope)^43^ in proliferating *S. pombe* cells (30°C). As shown in Fig. 5D, the lateral diffusion rate of both markers is reduced in *oca3Δ* cells, but importantly, the diffusion rate is recovered to *wild-type* values by addition of tween 20 (0,04%) in these mutant cells. Thus, the fluidizing properties of this non-ionic detergent most likely rescue growth defects in Oca3/Emc2-depleted cells by *in vivo* counteracting the rigidizing effects of excess ergosterol.

Under proliferating growth conditions (30°C), Oca3/Emc2-depleted cells aggregate the mitochondria (Fig. 2D-E) and decrease levels of mtDNA molecules (Fig. 2B). In addition to cold-sensitive growth, we show that tween 20 significantly rescues the defective mitochondrial tubular network (Fig. 5E) and mtDNA content (Fig. 5F). Overall, our results indicate that non-optimal membrane fluidity is an important biophysical contributor to the pleiotropic defects observed in Oca3/Emc2-depleted cells. Furthermore, this result underlines a major role of the EMC complex in membrane fluidity homeostasis by regulating ergosterol levels in *S. pombe* cells through Ltc1 biogenesis.

### Membrane rigidity hinders folding of ER membrane proteins

How fluidization of cell membranes with surplus ergosterol rescues cold-sensitive growth and mitochondrial dysfunctions in EMC-deficient cells is intriguing. EMC is expected to assist folding at the ER of a still incomplete set of ER membrane proteins.^5,9,19,44^ To estimate the levels of ER-unfolded proteins in EMC-deficient cells, we monitored fluorescence intensity in proliferating cells (30°C) expressing the ER-reporter mCherry-ADEL under the promoter of the BiP1/GRP78 protein, an abundantly expressed Hsp70 family member specifically induced by the unfolded protein response (UPR).^45–47^ As shown in Fig. 6A, basal levels of mCherry ADEL expression decorate the ER-membranes of *wild-type* cells. In contrast, BiP1-driven expression of mCherry-ADEL is greatly induced in Oca3/Emc2-depleted cells. The UPR senses the protein folding capacity of the ER^45^, indicating that EMC dysfunction severely provokes the accumulation of ER-unfolded proteins in *S. pombe* cells (ER stressed cells).

**Fig. 6.**
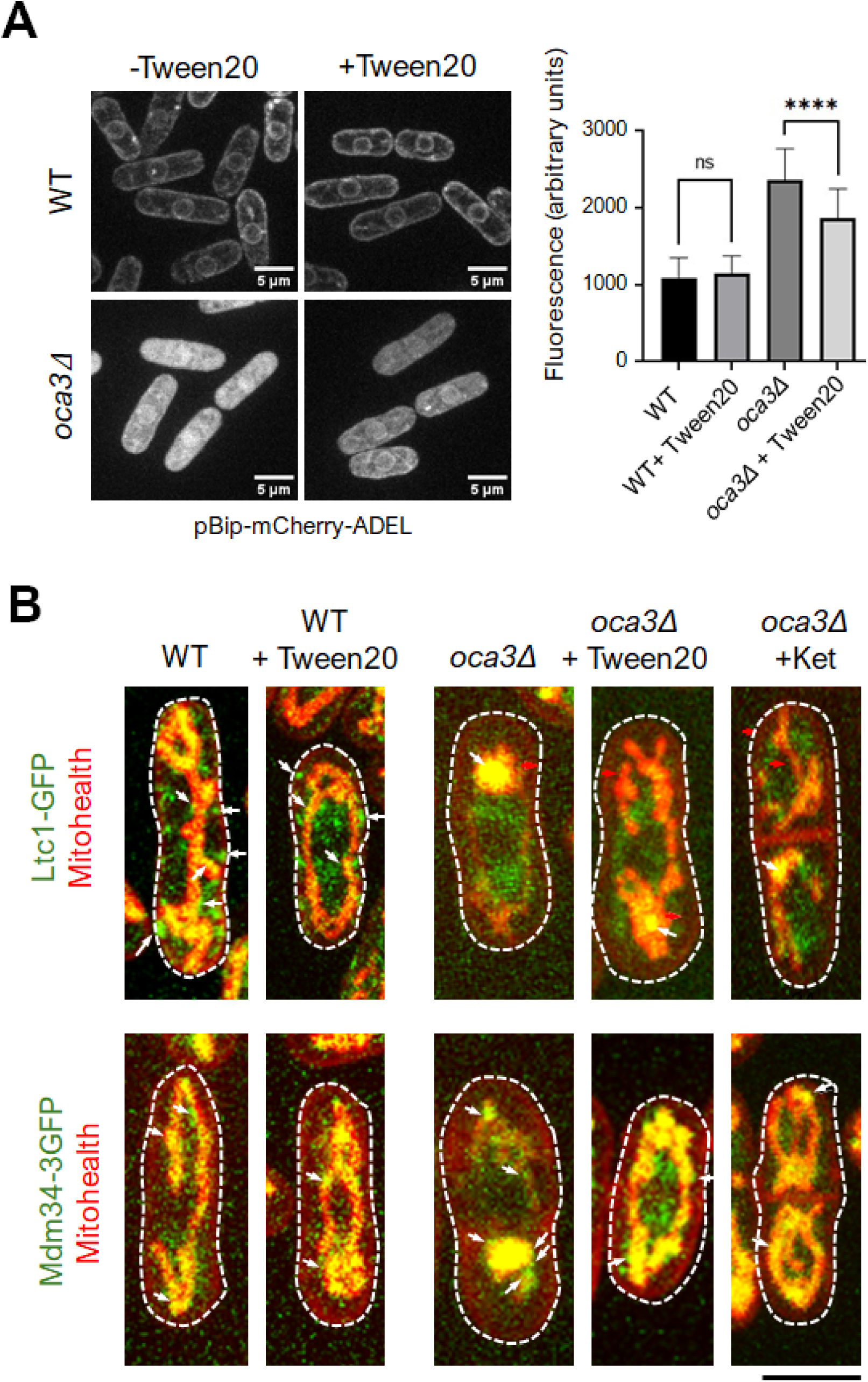
The UPR and ERMES and Ltc1 subcellular localization. **A.** mCherry-ADEL expressed under the Bip1 promoter in *wild-type* (wt, n=44) and *oca3Δ* (n=53) cells at 30°C (microphotographs) with or without tween 20 as indicated. Fluorescence quantification of mCherry-ADEL was used to estimate UPR levels. **B.** Representative proliferating cells (30°C) showing fluorescence images of Ltc1-GFP (upper panels) and Mdm34-3GFP (lower panels) in *wild-type* (wt) and oca3Δ cells in the presence and absence of tween 20 or ketoconazone as indicated. Mitohealth (red) and GFP (green) fluorescence are used in the same cells (merge images) to highlights mitochondrial and PM membrane localization and co-localization (yellow). Scale bar 5 µm.

When adding tween 20, a fraction of the UPR signal remains in these EMC-deficient cells, underlying a set of EMC clients which requires direct assistance of this complex for proper folding (see in Fig. 6A). But strikingly, tween 20 reduced BiP1-driven expression of mCherry-ADEL by approximately 20%. Biogenesis of transmembrane proteins is modulated by membrane lipid composition.^48^ We therefore conclude that in the absence of EMC function, ergosterol rigidized membranes impair proper folding of a significant fraction of membrane proteins at the ER, which is permitted (spontaneously or assisted by other chaperones) under optimal membrane fluidity conditions. Furthermore, our results give EMC a key role in membrane fluidity homeostasis to favor membrane proteins folding.

In view of this result, we analyzed the localization of Ltc1-GFP and Mdm34-3GFP in *oca3*Δ cells upon addition of tween 20. As shown above (Fig. 6B), this non-ionic detergent significantly rescues defects in the tubular structure of mitochondria, but Ltc1-GFP remains depleted from ER-PM and ER-Mitochondrial contact sites (Fig. 6B and Suppl. Fig. S2). Accordingly, ergosterol level remains high in tween 20 treated *oca3*Δ cells /Fig 5C). Therefore, Ltc1 requires EMC activity independently of the membrane fluidity, and recovery of the mitochondrial structure cannot be directly attributed to a tethering role of Ltc1. However, Mdm34-3GFP expands as the tubular structure of mitochondria is recovered in the presence of tween 20 (see in Fig. 6A and Suppl Fig. S1). The same results were obtained by ketoconazole treatment of *oca3*Δ cells (see in Fig. 6B and Suppl Fig. S2). These observations lead us to conclude that EMC assists membrane proteins folding and insertion by a direct action on client proteins (i.e. Ltc1), but also may facilitate the biogenesis of some other membrane proteins by providing optimal membrane fluidity (i.e. ERMES components).

## DISCUSSION

### Pleiotropic outcomes of EMC disruption arise from membrane rigidizing effects of surplus ergosterol

The EMC complex assists the folding and insertion of TMD-containing proteins at the ER with specific topologies so that they carry out their activities^19^. Given the large proportion of genes encoding membrane proteins, a central role for EMC as a TMD chaperone and insertion factor may explain its high abundance, broad conservation, and pleiotropic phenotypes.^9,49^ Accordingly, we observed that *S. pombe* cells losing EMC function constitutively overaccumulate ergosterol (Fig. 4A-B) and show cold-sensitive growth (Fig. 1D, Fig. 2H and Fig. 4C) and severe mitochondrial dysfunctions (Fig. 2D-E and Fig. 3).

Interestingly, in the routinary ketoconazole resistance assay to confirm the excess of ergosterol^31^, we observed that this drug recues both cold-sensitivity of growth (Fig. 4C) and mitochondrial dysfunctions in EMC-deficient cells (see in Fig. 5E and 6B). To our surprise, the non-ionic detergent tween 20 equally rescues cold-sensitive growth and mitochondrial dysfunctions, as well as ER stress, in these mutant cells (Fig. 6A-B and Suppl. Fig. S2). In all eukaryotes, cholesterol/ergosterol homeostasis is part of the adaptive mechanisms that maintain optimal cell membrane fluidity under changing environments.^32,33^ Decreased levels of EMC clients may impact in their functions in EMC-deficient cells.^9^ Nonetheless, in *S. pombe* EMC-deficient cells, we demonstrate that the pleiotropic phenotypic defects observed in EMC-deficient cells mainly arise from non-optimal membrane fluidity caused by the excess of ergosterol.

### EMC drains membrane ergosterol *via* the sterol transfer protein Ltc1

Sterols serve for membrane integrity and proper activity of multiple membrane proteins^50^, but importantly, sterol homeostasis is essential to attain optimal fluidity of cell membranes.^51,52^ In mammals, a key step in this system involves the transport of sterols from the PM to the ER and from the ER to the mitochondria, where it is metabolized to 27-hydroxycholesterol that rapidly exits the cell.^33,53–55^ Disruption of this flow pathway drives to the overaccumulation of this sterol upstream of the ER-Mitochondrial transit step.^56^

In *S. pombe*, the sterol transfer Ltc1 plays a major role in the ergosterol flux across ER-PM contact sites, where it resides^22^ (Fig. 3 and Fig. 6B). Here we show that Ltc1 also localises to ER-Mitochondrial membrane contacts (Fig. 3 and Fig. 6B). Thus, as previously reported in *S. cerevisiae* cells^23^, this protein may also facilitate ergosterol transfer from the ER to the mitochondria. Since Ltc1 depletion leads to ergosterol overaccumulation in *S. pombe* cells^22^, it is likely that sterol transport from the PM to the ER and from the ER to the mitochondria by this protein is required to drain ergosterol excess in these cells.

It is noteworthy that the localization of Ltc1 in contact sites between the ER and the PM and between the ER and the mitochondria depends on the activity of the EMC (see in Fig. 3 and Fig. 6B). Thus, in *S. pombe* cells, the role of EMC in ergosterol homeostasis, and in turn in membrane fluidity, is driven by proper folding and insertion of Ltc1, perhaps draining sterol by a pathway similar to that found in mammal cells.^33,53–55^ The recent finding of a tripartite membrane contact site between the PM, ER, and mitochondria^57^ may facilitate sterols flow from the PM to the ER and then from the ER to mitochondria.

### Unfolded Ltc1 delivery into the mitochondria

During the biogenesis of TMD-membrane proteins, unfolded protein species eventually arise during EMC maturation that require clearance from the ER membrane.^9,57^ EMC components show close interaction with the ERAD machinery.^8,58^ Accordingly, quantitative proteomic methodologies identified putative EMC clients, including some of its own subunits, by significant depletion in EMC-deficient cells (i.e. EMC3 in Fig. 1C).^5,44^ However, other membrane proteins in these cells are not correctly folded but not degraded.^49^ This set of non-degraded EMC clients likely uses ERAMS (ER-associated mitochondrial sequestration) rather than ERAD to clear unfolded proteins.^27^ As described here, the sterol transfer protein Lam6/Ltc1 belongs to this set of proteins that, in the absence of EMC, is delivered to the mitochondrial matrix for the sequestration of unfolded proteins. Unfolded proteins may also come from membrane-associated proteins that requires optimal membrane fluidity for proper folding (i.e. Mdm34) (Fig. 3 and Fig. 6B). In agreement with the high UPR signal in EMC-deficient cells (ER stress) (Fig. 6A), we propose that accumulation of these unfolded proteins provoke mitochondria stress and swollen it to the point that ruptures seams at ER-mitochondrial junctions (Fig. 2D, Fig. 3 and Fig. 6B). Membrane fluidization by tween 20 in these cells facilitates folding and relaxes mitochondrial stress enough to recover the tubular structure and ER-mitochondrial binding factors (mainly ERMES). ER-mitochondria contacts couple mtDNA synthesis with mitochondrial division in human cells.^59^ The loss of mtDNA in EMC-deficient cells and its partial recovery with tween 20 may also respond to loss and recovery of ER-Mitochondrial contact sites in *S. pombe* cells (Fig. 2B and Fig. 5F respectively).

Whether the loss of the tubular mitochondrial network in EMC-deficient cells is due to the role of EMC on the tethering functions of Ltc1 and/or ERMES, or the consequence of the tension between the ER and the mitochondrial provoked by the accumulation of unfolded proteins at the mitochondrial matrix (or both) remains an open question. But importantly, membrane fluidification significantly recovers both proteostasis and ER-Mitochondria tethering in EMC-defective cells (Fig. 6), cellular dysfunctions often associated to ageing and neurodegeneration.^60–63^

Cells following membrane fluidification by tween 20 reduce by approximately 20% the UPR signal due to EMC inactivation (Fig. 6). Multiple genetic studies in patients with congenital malformations have now suggested that variants in EMC subunits may be at the root of various congenital malformations.^3^ Changes in membrane fluidity in these variants with concomitant effects on membrane protein activities often accompany the transition from a healthy to a pathological state.^64^ Although deliberate modulation of membrane fluidity with drugs has not been exploited to date, our data suggest that the fluidizing effect of tween 20 (food additive E432), which counteract sterols rigidity in EMC-deficient cells, challenges conventional approaches to overcome proteostasis-related human defects and may open new insight into potential treatments related to neurodegenerative and EMC-associated diseases.

### Experimental Procedures

#### Media and growth conditions

Standard fission yeast growth media and molecular biology techniques were employed throughout the experiments as described in Moreno *et al*.^65^ Sporulation agar (SPA) was used for mating and sporulation. Tetrad pulling for segregation analyses was performed in a Singer MSM 400 automated dissection microscope (Singer Instruments). For spot tests, cells were cultured mid-log phase in Edinburgh Minimal Media 2 (EMM2) media. Cell number/mL was scored in Neubaueŕs chamber and matched dilutions were calculated for all cultures. Serial five-fold dilutions were plated onto solid media and incubated at the indicated temperatures. Drugs were added to both solid and liquid media in at the specified in-text indicated concentrations for sensitivity/resistance assays (nystatin, Sigma N6261; ketoconazole, Sigma 420600; N-Acetyl-L-cysteine, Sigma A9165; tween20, Sigma P1379). When growing at restrictive temperatures, cells were incubated in a water bath at 20 °C for 6 hours unless stated otherwise.

For the fluidization viability spot test (Fig. 6A), three concentrations were used for each compound, the indicated, half and double. Linoleic Acid (0.5mM, Sigma L1376), Sorbitol (1.5M, PanReac A4992), Ethanol (6% v/v, PanReac 131086), PEG 4000 (5 wt%, VWR 26606), tween20 (0.02% v/v, Sigma P1379) SDS (0.02% v/v, Sigma L3771), Triton X100 (0.02% v/v, Sigma X100) and Nonidet P40 (0.02% v/v, USB 19628). All compounds were added to plates of rich media (YES). Curated data from S. pombe genes were obtained from Pombase.^11^

#### Gene Tagging and genetics analysis

Gene tagging was performed according to described protocols^66^, using the pF6aMX6 plasmid series. Following transformation, PCR from target loci was employed to verify the integration of the tags at the expected genomic locations in the candidate strains. Tetrad analysis for single integrations and double mutant’s constructs was performed by tetrads dissection following described methods.^65^ Strains used in this study are listed in the Supplementary Table S1.

#### Fluorescence Microscopy

For live-cell imaging, cells were mounted in an Ibidi µ-Slide 8 Well chamber (Ibidi 80826) adhered to the chamber with soybean lectin (Sigma L1395). (Cells were kept alive in EMM2). Imaging was conducted using a spinning-disk confocal microscope (IX-81, Olympus; CoolSNAP HQ2 camera, Plan Apochromat 100x, 1.4 NA objective, Roper Scientific) and Metamorph software, with the temperature maintained at 20 or 30°C. For DAPI/Calcofluor Staining, we followed the protocol described in Hoyos-Manchado *et al*.^67^ Images were captured using a Nikon ECLIPSE Ti-S microscope equipped with a Plan Apo VC 60x/1.40 Oil NA lens.

All images were processed using the open-source software ImageJ (Fiji).^68^ Unless otherwise stated in the figure legend, the images presented in the figures correspond to maximal projections of 26 slices with a Z-step of 0.3 μm.

For MitoTracker-Hoechst staining in *S. pombe* cultures, cells were cultured in minimal medium until reaching an exponential phase of 2×10^6^cell/mL. MitoTracker CMxRos (200 nM in medium from the stock of 1 mM in DMSO, ThermoFisher M7512) and Hoechst (0.5-1 µg/ml in water from stock stored at 4°C, Sigma 63493) solutions were used for staining the cells.

Fluorescence Recovery After Photobleaching (FRAP) was performed using a spinning-disk confocal microscope (IX-81, Olympus; CoolSNAP HQ2 camera, Plan Apochromat 100x, 1.4 NA objective, Roper Scientific) and Metamorph software. A region of the cell was targeted, and recovery of the fluorescence was measured. Cells were incubated 30 minutes with cycloheximide (150 µm/mL, Sigma C7698) prior imaging.

#### mtDNA qPCR

To extract mitochondrial DNA (mtDNA) from *S. pombe*, cells were cultured in rich medium (YES) until exponential phase (about 2×10^6^ cell/mL) at the desired temperature (20 or 30 °C). Extract DNA from cultures using FastPrep-24™ 5G (Fisher Scientific) machine and later purification by phenol-chloroform. DNA concentrations were measured using a NanoDrop 2000 (Thermo Fisher Scientific) and adjusted to 20 ng/µL. Sample integrity was verified using a simple PCR with one of the qPCR primers. To perform the qPCR, 20 ng of DNA from the samples was added per triplicate, along with SYBR Green (Takara Bio RR820L) following the manufacturer’s instructions and 1 µL of each primer (Table S1) diluted to 100 nM. The qPCR was carried out with the following parameters: 1x 95°C for 3’ and 40x 95°C for 10” + 62°C for 30”, followed by a melting curve from 62°C to 95°C. The 2^-ΔCt^ and 2^ΔΔCt^ were calculated and the obtained data were normalized with respect to the *wild-type*

#### TLC of Ergosterol

Lipids were extracted from cells following the protocol by Markgraf *et al*.^69^ Briefly, cells were cultured until the exponential phase. Cells were collected and resuspended in 330 µL of methanol with glass beads and pulsed in a fast-prep. Then 660 µL of chloroform were added, centrifuged and the supernatant was recovered and 0.2 volumes of 0.9% NaCl were added. After the formation of two phases, the lower phase was recovered and evaporated completely using a SpeedVac. The residue was resuspended in 20 µl of chloroform. To perform lipid separation, TLC silica gel 60 F₂₅₄ plates of 10×20 cm were prepared, and samples of known concentrations (0.25, 0.5, and 1 mg/mL) of ergosterol (sigma E6510) were prepared. 10 µl of each sample and of the ergosterol solutions were deposited, leaving at least 1 cm of separation between them. The separation was performed in three phases following described protocols^70^ In summary, a first stage up to 50% of the maximum migration with hexane:diethyl ether:glacial acetic acid (50:50:2), a second stage up to 80% of the maximum migration with hexane:diethyl ether:glacial acetic acid (80:20:2), and finally a third stage to the maximum migration with 100% hexane. To reveal the lipids once separated, a ferric chloride solution (50 mg of FeCl_3_ · 6H_2_O in 90 mL of water) with 5 ml of glacial acetic acid and 5 mL of concentrated H_2_SO_4_ is sprayed over the TLC plate and heated at 100°C for 3 minutes.^71^ Plates were scanned using a GelDoc Go (BioRad 12009077) system and quantified in ImageJ software using the standard curve of pure ergosterol.

#### Statistical Analysis

Graphs and statistical analyses were produced using Microsoft Excel and Prism 9.0 (GraphPad Software). The graphs display the mean and standard deviation (SD), with the number of cells analyzed (n) indicated from at least three independent experiments. For comparisons between two groups, an unpaired Student’s t-test was used. Multiple-group comparisons were conducted using non-parametric Kruskal-Wallis one-way ANOVA and ordinary one-way ANOVA.

## Acknowledgments

We thank Dr. Martin for providing D4H probe plasmid and *ltc1*-GFP-tagged strains and Dr. Daga for providing fluorescent tagged strains. This work was supported by the Spanish Ministerio de Ciencia e Innovación (grant numbers PID2019-111124GB-I00 to J.J.). We thank Katherina García for her assistance in the advanced microscopy facility, Victor Carranco for excellent technical assistance, and all members of the yeast genetics group at the CABD for valuable comments and discussions.

## Authors contributions

M.B., V.A.T. and J.J. designed the experiments. M.B. performed the experiments and analyzed the data. V.A.T. analyzed the data and supervised experimental protocols. J.J. analyzed the data, acquired funding and wrote the paper with input from M.B. and V.A.T.

