## Supplemental Figs S1, S2 and Table 1 for "EMC regulates cell membrane fluidity to facilitate biogenesis of membrane proteins"

### Supplementary material

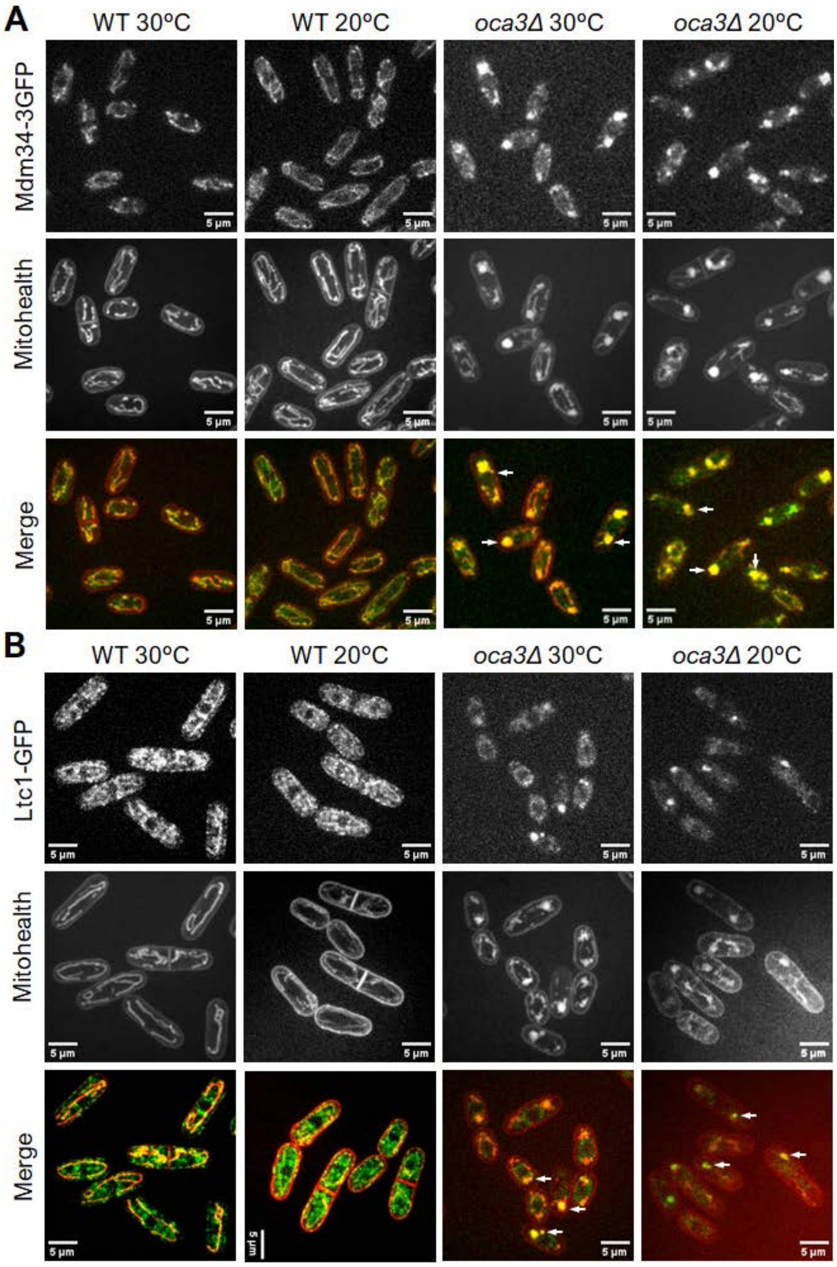


Suppl. Fig. S1. ERMES and Ltc1 subcellular localization A. Fluorescence images of the ERMES subunit Mdm34-3GFP in wild_type (wt) and oca9K cells at 20ºC and 30ºC (top panels). Mitohealth staining is used in the same cells (central panel) to highlights mitochondrial membrane localization (Merge, bottom panels). B. Fluorescence images of Ltc1-GFP in wild_type (wt) and oca9K cells at 20ºC and 30ºC (top panels). Ltc1-GFP localizes to cortical and internal dots (arrowheads) at the ER and Mitohealth staining is used within the same cells (central panel) to highlight mitochondrial membrane localization (Merge, bottom panels). Scale bar 5 µm.

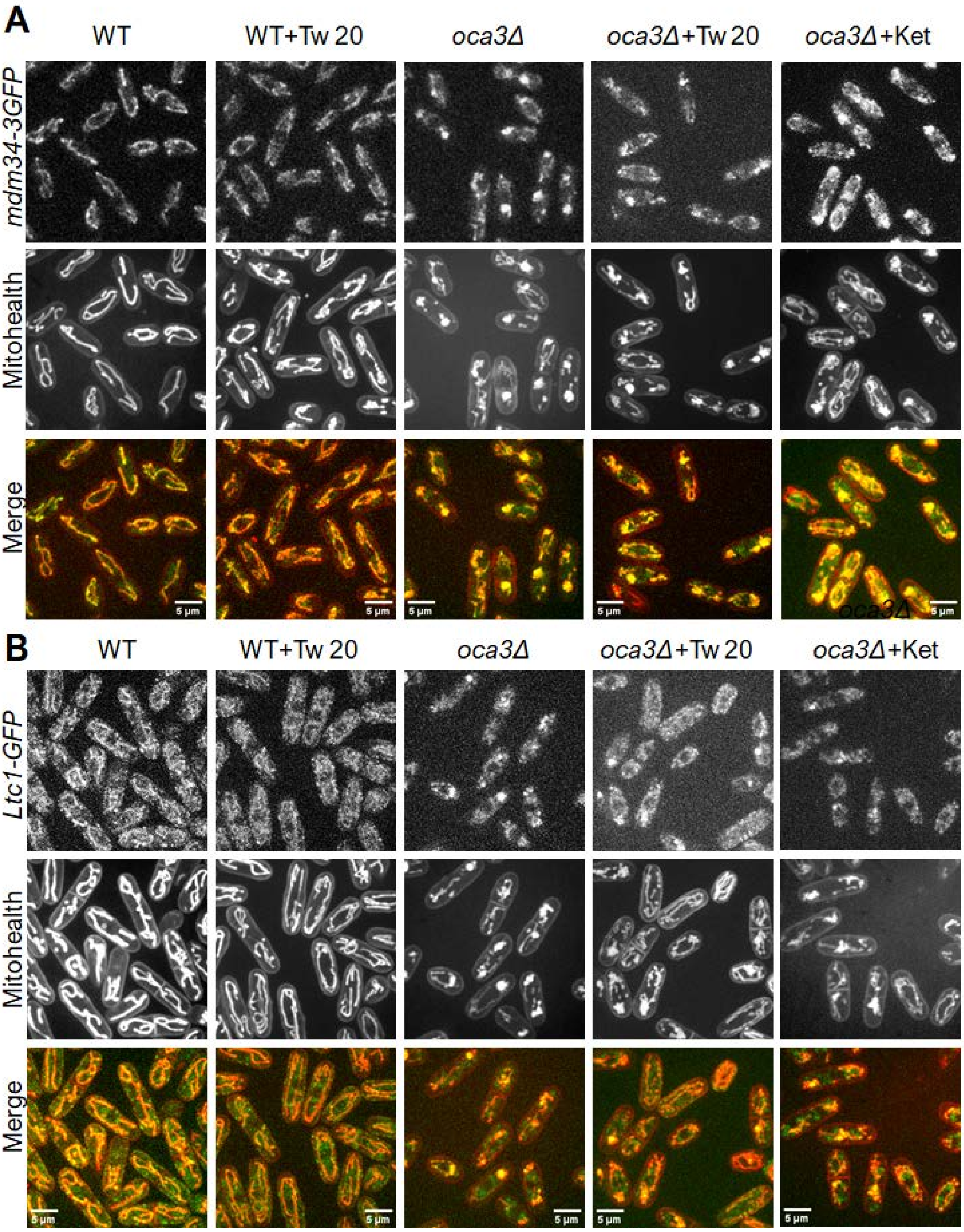


Suppl. Fig. S2. ERMES and Ltc1 subcellular localization A. Fluorescence images of the ERMES subunit Mdm34-3GFP in wild_type (wt) and oca9K cells at 20ºC and 30ºC (top panels). Mitohealth staining is used in the same cells (central panel) to highlights mitochondrial membrane localization (Merge, bottom panels). B. Fluorescence images of Ltc1-GFP in wild_type (wt) and oca9K cells at 20ºC and 30ºC (top panels). Ltc1-GFP localizes to cortical and internal dots (arrowheads) at the ER and Mitohealth staining is used within the same cells (central panel) to highlight mitochondrial membrane localization (Merge, bottom panels). Scale bar 5 μm.

Supplementary Table S1. Strains used in this study

| Strain | Source | Genotype |
| --- | --- | --- |

| | | |
|---|---|---|
| VA1 | VAT Collection | h- 972 |
| JJ1209 | JJ Collection | h- nda3-km311 ura4-d18 leu1-32 |
| JJ2364 | Bioneer | h+ oca3::KanMx4 leu1-32 ura4-d18 ade6-M210/216? |
| JJ2366 | This study | h+ Oca3:mcherry:hph ura4-D18 his2 leu1.32 |
| JJ2374 | This study | h+ oca3::Kan |
| JJ2375 | This study | h- oca3::Kan |
| JJ2408 | This study | h? oca3::kanMX4 ish1-GFP:kan |
| JJ2409 | R. Daga Lab | h- ish1-GFP::kan ura4.d18 |
| JJ2414 | This study | h? oca3::kanMX4 scs2-GFP:ura ura- |
| JJ2415 | R. Daga Lab | h? scs2-GFP:ura ura- |
| JJ2424 | This study | h- emc3::hph |
| JJ2426 | FY33691 | h+ coq4::kanMX6 leu1-32 ura4-D18 |
| JJ2453 | This study | h- emc3-tomato:hph |
| JJ2454 | This study | h- emc6-tomato:hph |
| JJ2468 | This study | h- oca3::kan emc6-tomato-hph |
| JJ2471 | This study | h+ oca3::kan emc3-tomato-hph |
| JJ2490 | This study | h- emc6::hph |
| JJ2492 | This study | h+ emc5::kan his- |
| JJ2493 | FY17341 | h- erg5::ura ura- leu- |
| JJ2507 | This study | h? emc5::kan emc6-tomato:hph |
| JJ2511 | This study | h? emc6::hph emc3-tomato:hph |
| JJ2512 | This study | h? emc5::kan emc3-tomato:hph |
| JJ2546 | Transformed VA1 from S. Martin plasmid PSM2056 | h- D4H-mCherry |
| JJ2547 | Transformed JJ2375 from S. Martin plasmid PSM2056 | h- oca3::kan D4H-mCherry |
| JJ2570 | This study | h? oca3::kan emc5-GFP:kan |
| JJ2571 | This study | h? emc5-GFP:kan emc6::hph |
| JJ2572 | This study | h+ emc5-GFP:kan |
| JJ2579 | This study | h? emc3::hph emc5-GFP:Kan |
| JJ2591 | This study | h- emc3::hph emc6-tomato:hph |
| JJ2606 | R. Daga Lab | h+ arg11-mCherry:nat |
| JJ2607 | This study | h- oca3::kanMX4 arg11-mCherry:nat |
| JJ2795 | This study | h- mdm34-3GFP:Kan |
| JJ2798 | This study | h- mdm34-td.tomato:hph oca3::kanMX4 |
| JJ2815 | This study | h? oca3:mcherry:hph ADEL-GFP:leu |
| JJ2818 | S. Martin Lab YSM3473 | h+ ltc1::hphMX6 ura4::pPom1-Ltc1-sfGFP::kanMX6 |
| JJ2820 | This study | h? oca3:mcherry:hph ura4::pPom1-Ltc1-sfGFP::kanMX6 ltc1-sfGFP::kanMX6 |
